## Supplemental Files for "SNAP-25, but not SNAP-23, is essential for photoreceptor function and survival in mice"

### Supplemental Information

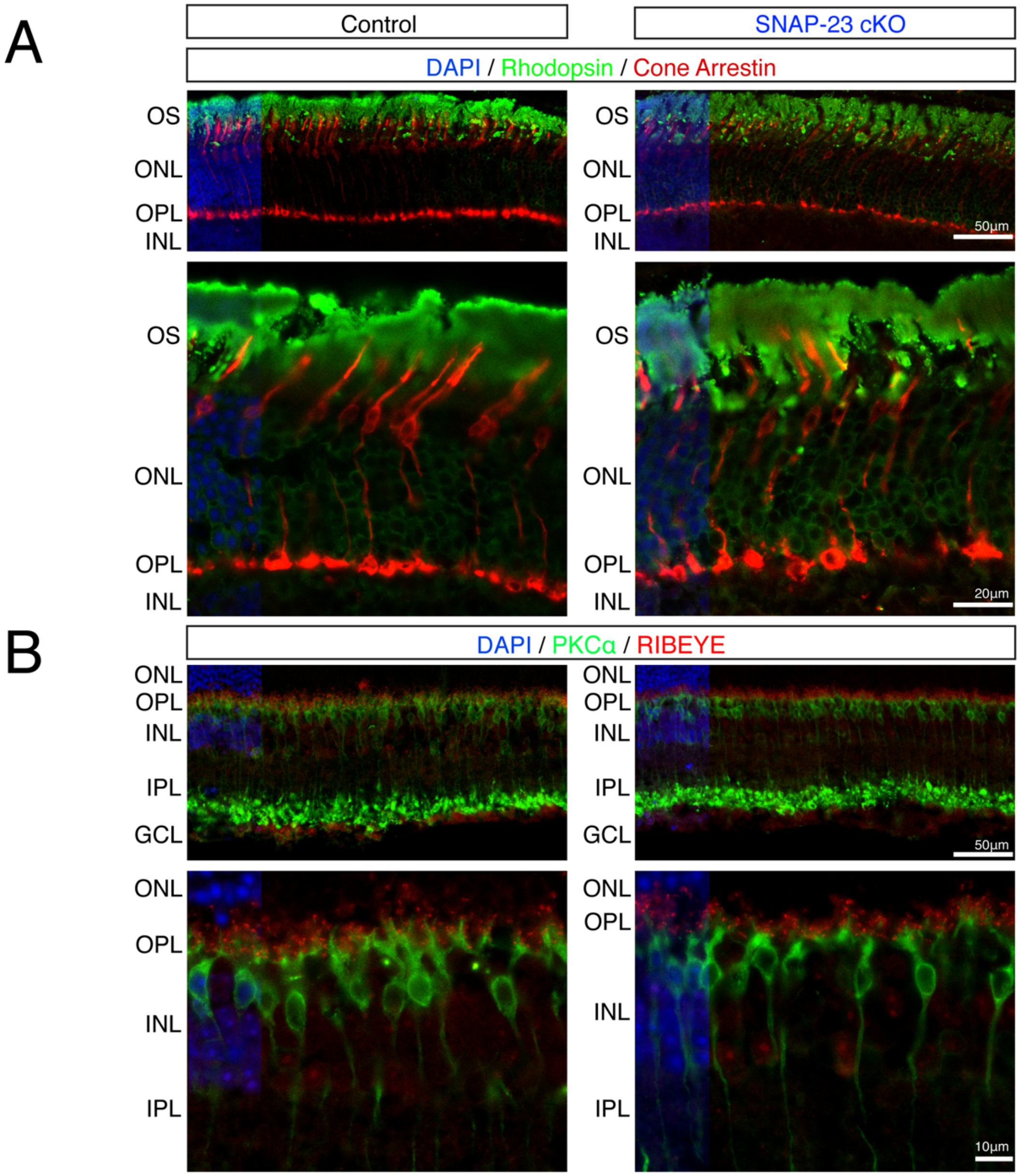


**Supplemental figure 1. Removal of SNAP-23 does not alter retinal integrity.**

(A) Immunostain of control and SNAP-23 cKO retinas for rods (green) and cones (red). Rods and cones are both present in SNAP-23 cKO retinas and stratify properly.
(B) Immunostain of control and SNAP-23 cKO retinas for rod bipolar cells (green) and synaptic ribbons (red). Synaptic ribbons are present in the presynaptic area of the synaptic area in both control and SNAP-23 cKO retinas. Synaptic ribbons form proper connections with post synaptic rod bipolar cells in both control and SNAP-23 cKO retinas.

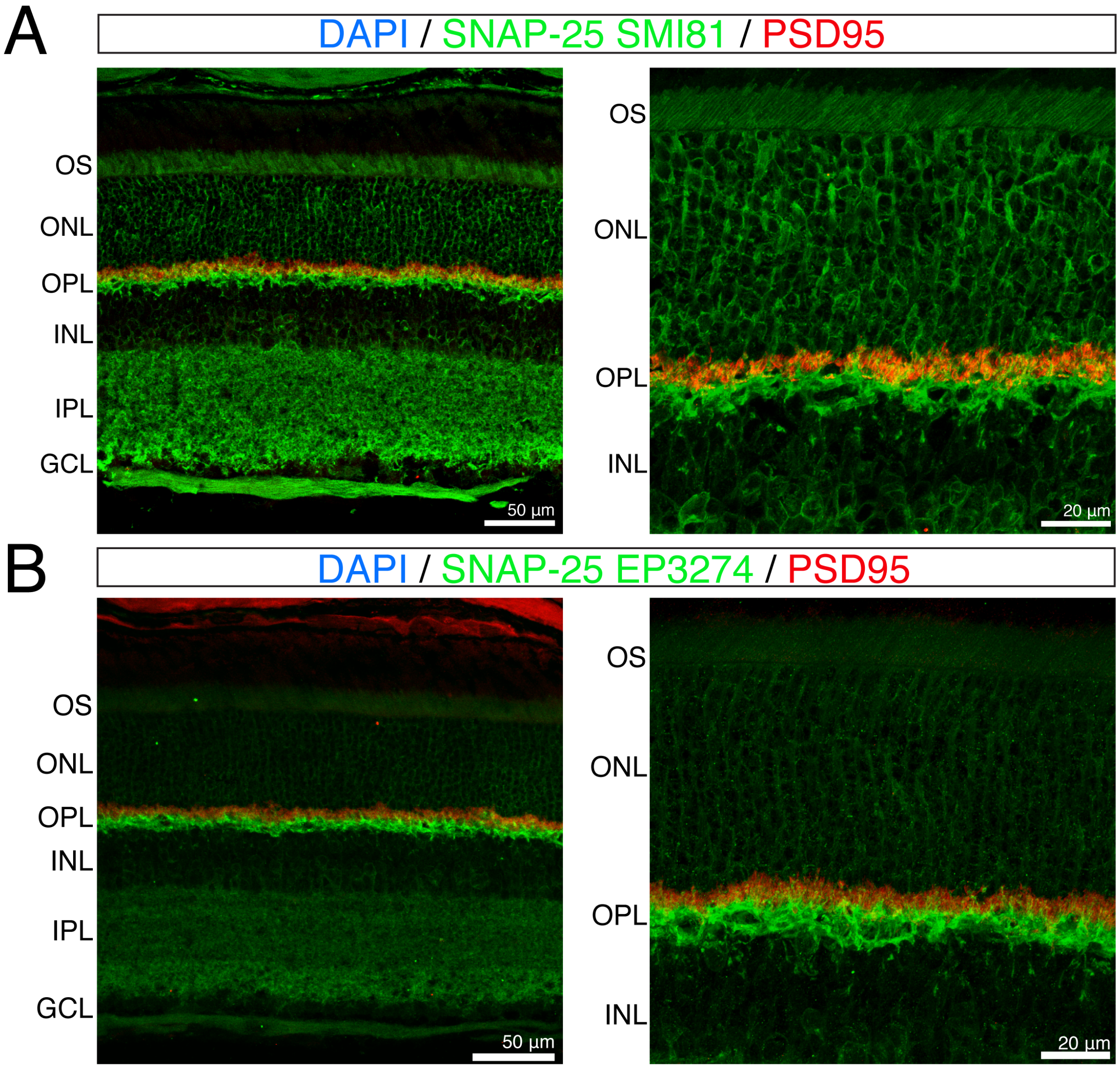


**Supplemental figure 2. SNAP-25 protein can be detected by antibody staining.**

(A) SNAP-25 SMI81 antibody staining (green) finds SNAP-25 throughout the retina, particularly in photoreceptor outer segments, surrounding photoreceptor nuclei, and in photoreceptor synaptic terminals. SNAP-25 colocalizes with presynaptic photoreceptor terminal marker PSD95 (red). SNAP-25 is also observed in the inner plexiform layer (IPL) and ganglion cell layer.
(B) SNAP-25 EP3274 antibody staining (green) finds similar SNAP-25 presence in photoreceptor as with the staining observed in (B).

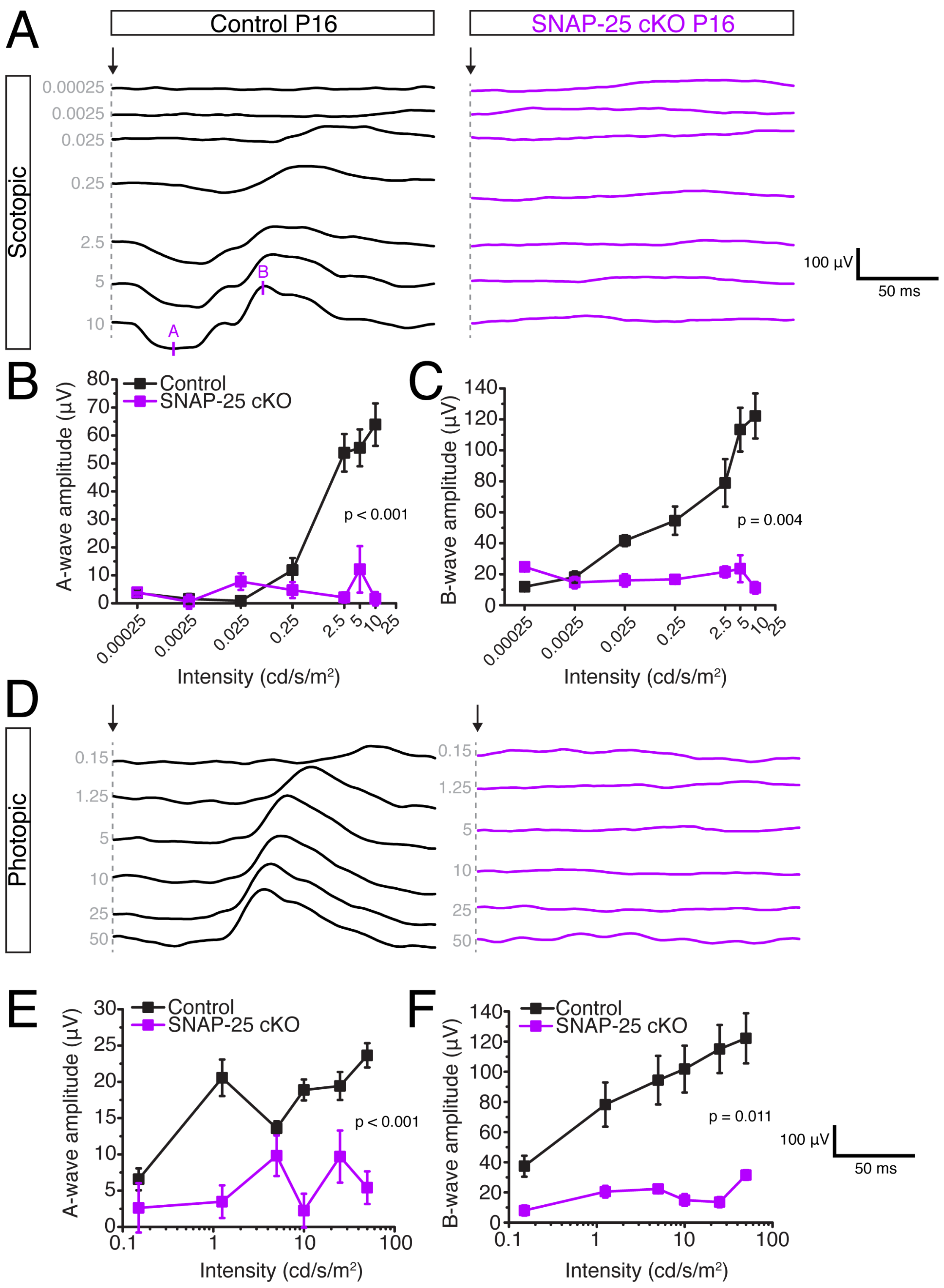


**Supplemental figure 3. No visual function in SNAP-25 conditional knockout mice at postnatal day 16.**

(A) Representative scotopic electroretinogram traces across 7 light intensities. Arrow indicate stimulation onset, * indicates b-wave. SNAP-25 cKO mice exhibit no responses to light stimulation at any light intensity at postnatal day 16.
(B) Quantifications of scotopic b-wave amplitudes. (n = 5 mice for both control and SNAP-25 cKO for all quantifications)
(C) Quantifications of scotopic time to peak of ERG b-wave.
(D) Representative photopic electroretinograms across 6 light intensities. Arrow indicates stimulation onset.
(E) Quantification of photopic b-wave amplitudes.
(F) Quantifications of photopic time to peak of ERG b-wave.

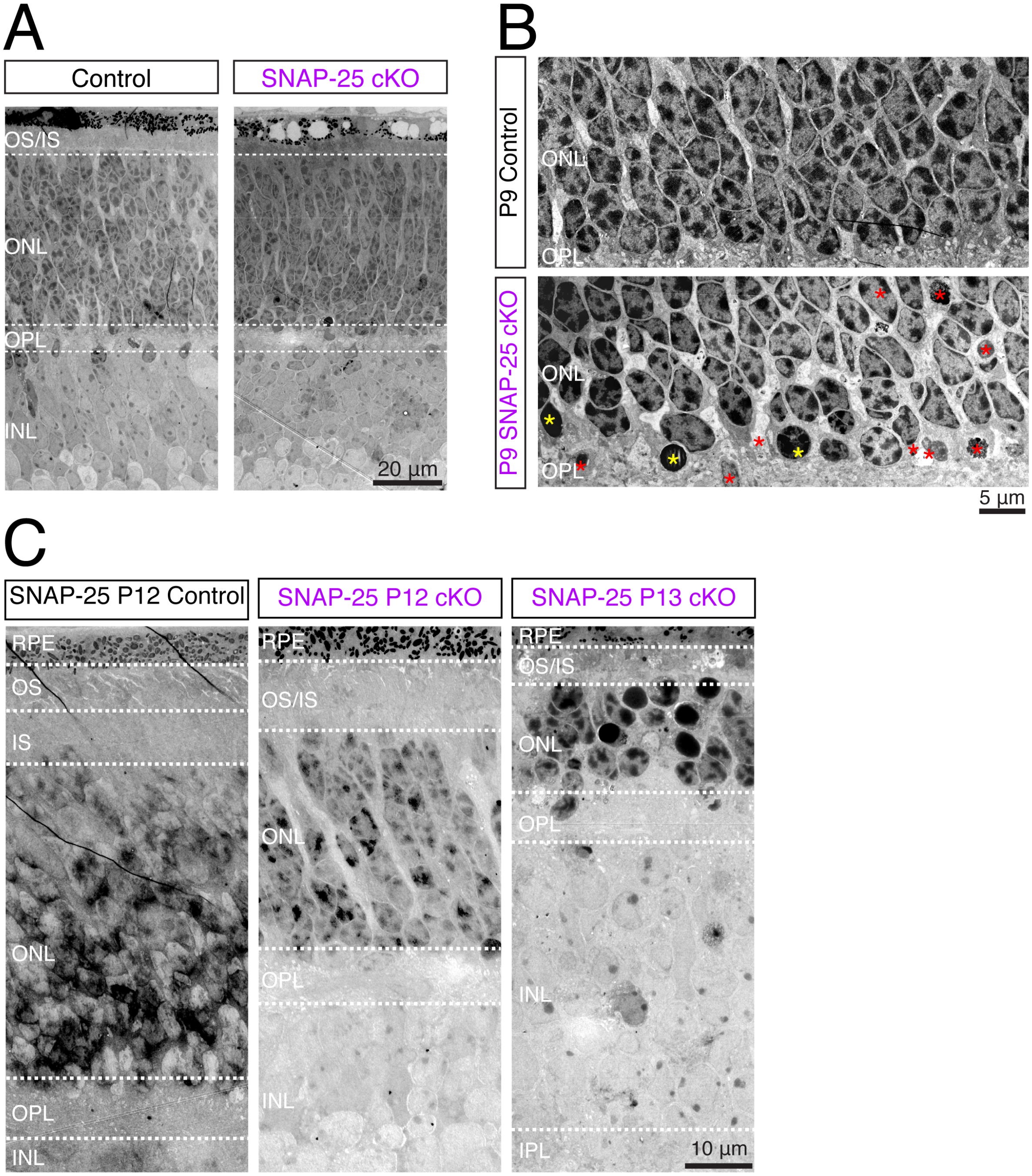


**Supplemental figure 4. Nuclear changes associated with the degeneration of photoreceptors**

(A) Retina thickness looks comparable between control and SNAP-25 cKO groups at postnatal day 9 (P9).
(B) Electron micrographs of control and SNAP-25 cKO retinas at postnatal day 9 (P9). At P9, photoreceptor nuclei exhibit conventional chromatin organization and have peripheral heterochromatin clusters. Chromatin structure is altered in unhealthy photoreceptors and uniformly dark condensed nuclei (yellow asterisks) can be observed in SNAP-25 cKO retinas. Other abnormal photoreceptors can be observed (red asterisks) and have features such as vacuolation, loss of membrane integrity, and fragmentation.
(C) Electron micrographs of control and SNAP-25 cKO retinas at P12 and SNAP-25 cKO retinas at P13. Very rapid degeneration of the outer nuclear layer occurs between P12 and P13.

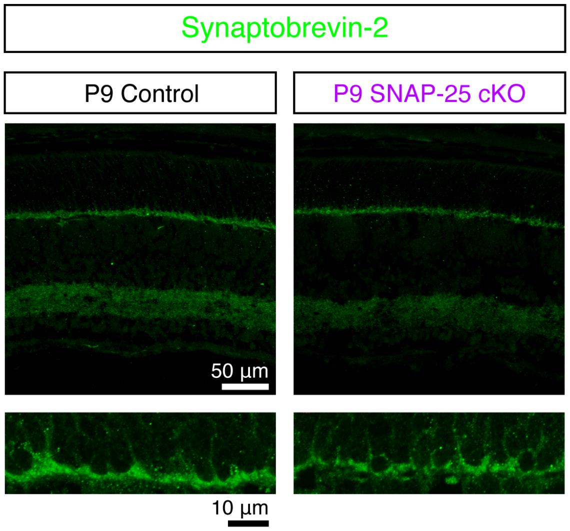


**Supplemental figure 5. Normal expression of synaptobrevin-2 in control and SNAP25 cKO retinas at P9.**

Immunostain of P9 control and SNAP-25 cKO retinas for synaptobrevin-2 (green). Synaptobrevin was observed presynaptically in photoreceptor terminals and in the inner plexiform layer.
